# Ancient stickleback genomes reveal the chronology of parallel adaptation

**DOI:** 10.1101/2025.04.14.648705

**Authors:** Jan Laine, Jana Nickel, Anders Romundset, Andrew D. Foote

## Abstract

Parallel evolution of traits and their underlying genetic basis is well-studied, however, studies of parallel chronology of adaptive genetic changes remain scarce. Threespine stickleback are a model system for studying parallel evolution. We present genomic data from nine subfossil stickleback bones dated to 14.8-0.7 KYR BP in age. Comparing the four highest coverage genomes, which represent different stages along the marine-freshwater continuum, we find that freshwater ancestry is often clustered rather than randomly distributed throughout the marine-freshwater divergent regions of the genome. We consistently find freshwater ancestry on chromosome IV at the early stages of freshwater adaptation. Regions of chromosome IV contain the greatest genetic differentiation between marine and freshwater ecotypes and among the highest density of quantitative trait loci. These include *EDA*, a large-effect pleiotropic locus associated with defensive armour and variation in neurosensory and behavioural traits. Freshwater ancestry is also found at inversions in the subfossils early in the adaptive process. Our findings add to the growing body of evidence that freshwater adaptation in threespine stickleback has a staggered but predictable temporal dynamic. The probability of parallel genetic change should be increased by the clustering of adaptive alleles in regions of suppressed recombination, particularly for genes that have a large phenotypic effect.

## Introduction

Genomic architecture is known to have an important effect on the use (and re-use) of genetic variation in adaptation (Westram et al. 2022). For example, the clustering of adaptive alleles in regions of suppressed recombination, particularly for genes that have a large phenotypic effect, can result in increased selection on these regions during ecotype formation (e.g. Huang et al. 2025; Meyer et al. 2024; Koch et al. 2022). Therefore, we predict that such genomic regions would show local adaptation at the early stages of ecology-driven selection. Parallel evolution offers a suitable paradigm to test this hypothesis.

Environments can impose re-occurring challenges and opportunities, which can drive parallel microevolutionary changes in allele or haplotype frequencies in homologous genes to promote similar adaptations among populations or species separated in space or time (Stern 2013). Different lineages can acquire the adaptive alleles independently through *de novo* mutations, or adaptive alleles can be already present in the ancestral population as standing genetic variation (SGV) (Martin and Orgogozo 2013). SGV can be more readily available for natural selection to act upon, thereby increasing the tempo of the adaption under strong directional selection (Barrett and Schluter 2008; Schluter and Conte 2009). Despite the increasing prevalence of studies of parallel evolution in nature (e.g. Bolnick et al. 2018; Stuart et al. 2017; Louis et al. 2021; Van Belleghem et al. 2018; Papadopulos et al. 2021), there is comparatively scarce data on parallelism of the chronology of adaptive genetic changes in these systems.

The threespine stickleback (*Gasterosteus aculeatus*) is considered a model organism in studies of parallel morphological and physiological adaptation underpinned by parallel genetic changes in response to ecological change (Reid et al. 2021). Recurrent adaptation of marine stickleback populations to freshwater lakes formed since the deglaciation has been well-documented as an exemplar case of parallel adaptive evolution (Bell & Foster 1994; Reid et al. 2021; Jones et al. 2012). Following the retreat of ice sheets at the end of the last ice age, isostatic crustal rebound has lifted bedrock depressions above sea level, causing a transition, from marine through a brackish water intermediate phase, to form freshwater lakes (Romundset et al. 2018). Stickleback populations in these newly formed lakes faced strong directional selection as they underwent this environmental transition. This led to the evolution of distinct marine and freshwater ecotypes across the Northern hemisphere range of the species (Bell and Foster 1994; Jones et al. 2012). These ecotypes differ in multiple traits, such as armour plate coverage, body shape, and salt handling (Bell 1976; Bell and Foster 1994). Key genetic loci underpinning parallel freshwater phenotypic divergence have previously been identified (Colosimo et al. 2005; Shapiro et al. 2004; Chan et al. 2010; Miller et al. 2014; Greenwood et al. 2016; Peichel & Marques 2017; Archambeault et al. 2020; Stepaniak et al. 2021). Colonizing freshwater populations are hypothesised to draw the beneficial freshwater alleles from the standing genetic variation present in the ancestral marine populations (Colosimo et al. 2005; Jones et al. 2012; Roberts Kingman et al. 2021). The ‘Transporter hypothesis’ suggests freshwater stickleback populations then act as reserves for the alleles crucial in the freshwater adaptation and replenish these alleles in the marine populations through low levels of gene flow from freshwater populations (Schluter and Conte 2009).

Analyses of the first genome sequences from stickleback sampled across the species’ Northern hemisphere range (Jones et al. 2012) identified 242 marine-freshwater divergent loci (*i*.*e*. those at which different alleles were associated with either marine or freshwater populations of threespine stickleback). Marine-freshwater divergent loci are unevenly distributed across the stickleback genome, clustering disproportionately on a subset of chromosomes (Jones et al. 2012; Roberts Kingman et al. 2021). Quantitative trait loci (QTL) within these regions vary in the effect size on the phenotypic traits they regulate (Peichel & Marques 2017; Schluter 2021; Liu et al. 2022). Many marine-freshwater divergent loci are assembled in close linkage to each other in regions of low recombination rate, creating units or “cassettes” that more likely segregate as one complex carrying a set of multiple clustered adaptive alleles (Jones et al. 2012; Liu et al. 2022; Venu et al. 2024). Finally, low recombination regions are flanked by recombination hotspots (Samuk et al. 2017; Roberts Kingman 2021; Venu et al. 2024). This is hypothesised to allow the freshwater associated alleles to be rapidly re-assembled in these cassettes in response to the selective pressures associated with freshwater environments, and to allow the dis-association between freshwater haplotypes by recombination in marine populations (Schluter and Conte 2009).

Previous studies utilizing time-series genomic data from capture-release experiments of marine fish into freshwater habitats, and ancient DNA (aDNA) data from the late Pleistocene bones and sediments, have shed light on the chronology and tempo of the early stages of the freshwater adaptation (Kirch et al. 2021; Roberts Kingman et al. 2021; Laine et al. 2024). Despite the differences in the study setting: namely gradual environmental transition from marine to freshwater environment (aDNA studies), compared to direct transfer of marine sticklebacks to freshwater lakes (experimental releases); both types of study found freshwater alleles at some of the same major effect loci shortly after the colonisation of freshwater (Kirch et al. 2021; Roberts Kingman et al. 2021; Laine et al. 2024). In this study, we further explore these dynamics utilizing four partial genomes acquired from an initial dataset of 13 ancient threespine stickleback samples collected from isolation lakes across the Norwegian coast and dating 13 – 0.7 KYR BP (Supplementary Table 1). The bone samples were collected from sediment layers representing the transitional brackish water environment, where the early onset of the directional selection for the freshwater phenotype would have taken place. We compare freshwater ancestry across the ancient stickleback genomes to published genomes from recent capture-release experiments (Roberts Kingman et al. 2021; Supplementary Table 2). Our goal is to understand the temporal dynamics, specifically, the predictability of the chronology of the detectable effects of selection on freshwater-associated alleles. We predict that if features of the genomic architecture such as localised reduced recombination and strong linkage play a role in promoting freshwater adaptation, then we will find parallelism in the chronology of genetic changes across the marine-freshwater divergent regions of the genome.

## Materials and Methods

### Ancient sample collection

Ancient stickleback bones were collected from sediment cores retrieved with either a ‘‘Russian-type’’ peat corer or percussion coring systems depending on the lake depth during the years 2019-2022 (Figure 1B, Supplementary Table 1; Jowsey 1966; Reasoner 1993; Laine et al. 2024). Sediment cores were transported to the laboratory at the Geological Survey of Norway (NGU), where 1 cm thick slices were wet-sieved through 125 µm sieving membrane, and residuals inspected for bone findings under a light microscope. All sub-fossils were found in transitional brackish water layers except for lake K-18, where stickleback bones were found in two separate marine layers 150 cm apart (depths of 840 cm and 990 cm). Some of these lakes had two distinct freshwater transitions, as temporarily increased sea levels had re-connected them back to the marine fjord after the initial isolation. Whether the subfossils were found from the first or second transition is indicated in the Supplementary Table 1. All ancient hard parts recovered from sediment cores were kept at -80ºC until the DNA extraction step. See Supplementary Figure 1 for stratigraphy plots.

**Figure 1.**
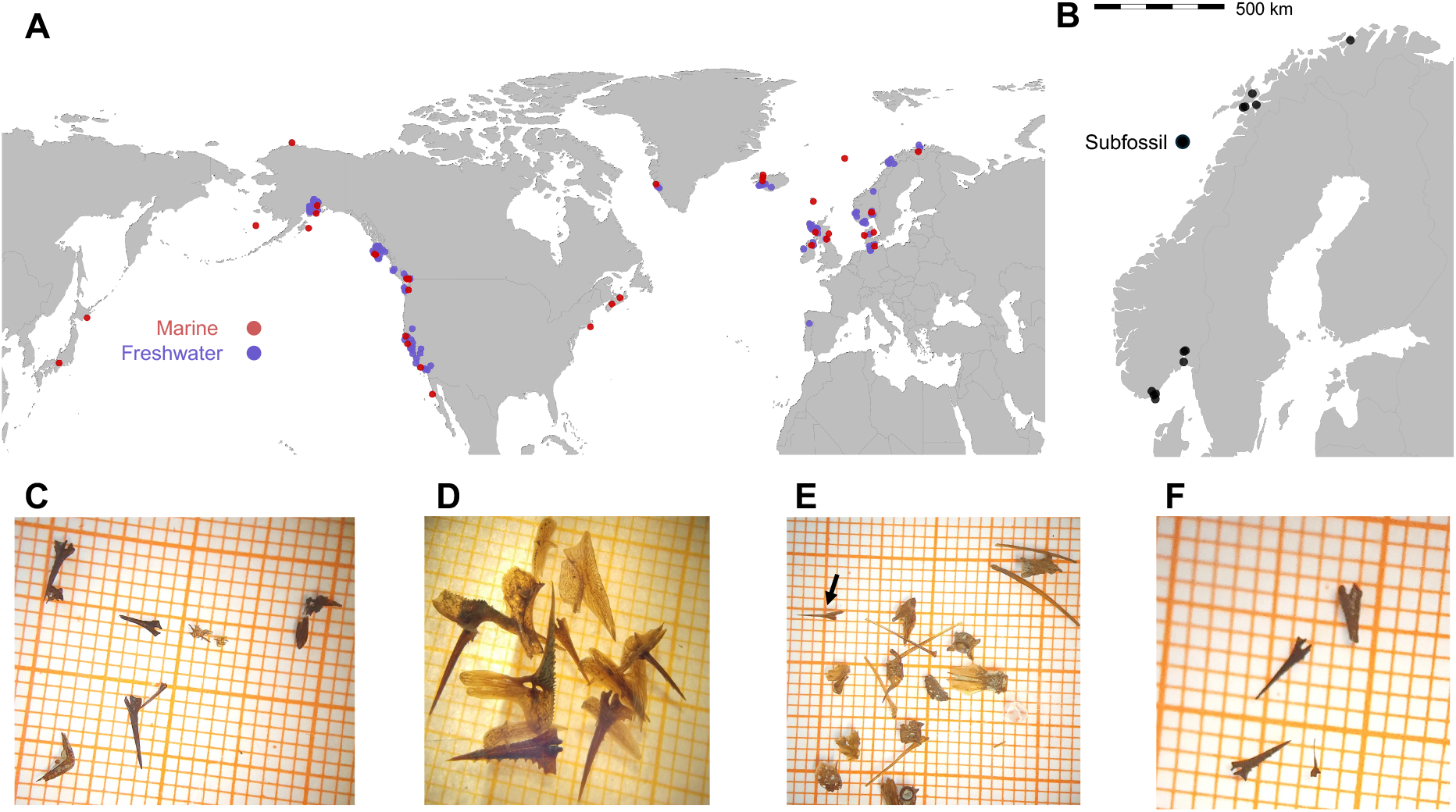
**A**. Map of the sampling locations of the 553 contemporary threespine stickleback included in this study. Marker colours denote the ecotype. **B**. Map of the locations where ancient stickleback bones, spines and armour plates included in this study were found in sediment cores by the Norwegian Geological Survey. **C-F**. Examples of ancient stickleback remains included in this study. The samples were found in cores from the following lakes: **C**. Ravnsborgtjernet (10.3 KYR BP); **D**. Jossavannet (12.9 KYR BP); **E**. Lake K18 (14.8 KYR BP). Note that many of the bones in panel **E** were genetically identified as sculpin – which can be a predator of threespine stickleback (McPhail 2007; Reimchen 1994; but see Roesti et al. 2023), the black arrow indicates a threespine stickleback spine; **F**. Dingneset (0.7 KYR BP).

### Ancient DNA extraction

Ancient DNA lab work was conducted in the dedicated ancient DNA facilities of the University of Copenhagen and NTNU. Details for the extraction of DNA from ancient stickleback bones and spine from the Jossavannet and Klubbvatnet samples are given in Kirch et al. (2021) and Laine et al. (2024). Bones from two lakes, Hoksvannet and Borrevannet, were treated as per Kirch et al. (2021); Bones from lake Lilletjønn were treated as per Laine et al. (2024). Ancient bone samples from eight further lakes were treated as per Laine et al. (2024), but with an additional extended lysis step.

Briefly, DNA was extracted using a silica-based method, where each individual bone or spine was incubated overnight under motion at 55°C in 500 µl extraction buffer (0.45M EDTA, 0.1M Tween 20, 100 mg proteinase K). Each sample was then centrifuged at 2300 rpm for 5 min and the supernatant was collected and concentrated and purified using a QIAGEN MinElute spin column (QIAGEN, Valencia,CA, USA). Remaining bone material after the first lysis were treated with an additional lysis step with an extended 72-hour incubation.

### Library preparation

Double-stranded Illumina libraries had been previously constructed using the blunt-end single tube (B.E.S.T.) method of Carøe et al. (2018) for the Jossavannet and Klubbvatnet samples and those libraries were then sequenced to saturation for the Kirch et al. study (2021). The same double stranded library build approach was used for the Borrevannet and Hoksvannet samples. Single-stranded DNA libraries were constructed using the method in Kapp et al. (2021) and 20 µl of input DNA extract per library. Extracts were those from the Kirch et al. (2021) study and all other ancient bone extracts generated for this study. The adapters and single-stranded binding proteins (SSB) were prepared and diluted following the recommendation for concentration defined by the DNA input tier system described in the protocol published by Kapp et al. (2021). As a conservative approach to avoid adapter dimer, we used tier 5 in all reactions. Libraries were purified with multiple ethanol washes whilst being retained on biotinylated beads and subsequently eluted in 35 µl of modified Qiagen elution buffer (containing 0.05 % Tween-20). A volume of 15 µl of DNA library template was used for dual-indexing PCR amplification following a AmpliTaq Gold (ThermoFisher Scientific) protocol at 25 cycles for each sample. Each 50 µl PCR reaction contained 15 µl of library, 1 µl of AmpliTaq Gold polymerase, 1 µl of 50 mM dNTP, 5 µl of 10x PCR buffer, 1 µl of 125 mM MgCl_2_, 0.4 µl of Bovine Serum Albumin (BSA), 1 µl of both 10 µM Illumina P5 and P7 index primers and was made up to 50 µl with molecular grade water. PCR temperature profile included an activation step at 95°C for 5 min, followed by 25 cycles of denaturation at 95°C for 30 s, annealing at 55°C for 30 s and elongation at 72°C for 1 min, with a final extension step at 72°C for 7 min. The final PCR product was purified in two consecutive treatments of NEBNext^®^ Sample Purification Beads (E7659) in 1.2x ratio to the library pool. Extraction, library build and index PCR blanks were included to evaluate potential contamination during the library building process. Multiple PCRs were performed on each library to maximize complexity for downstream analyses.

### Target enrichment capture

To maximize the coverage at marine-freshwater informative sites in the genome, we performed target capture enrichment experiments using biotinylated RNA baits (Enk et al. 2014; MyBaits, Daciel Arbor Biosciences, Design ID: D1029010KNT, Ref#: 220126-900, see Laine et al. 2024). The 38,391 biotinylated RNA baits targeted 10,026 transversions in regions of the genome previously identified as being consistently differentiated between marine and freshwater populations in a global dataset (Jones et al. 2012), see Laine et al. (2024) for more details on target site coordinates and individual bait sequences.

Amplified libraries were subject to enrichment following the manufacturer’s High Sensitivity protocol, but with a single hybridization step at 60°C and no post-capture PCR. Enriched and non-enriched libraries were then sequenced across a lane of an Illumina NovaSeq 6000 S4 flow cell. All raw sequence data are available at NCBI Bioproject ‘Stickle-back-in-time’ (PRJNA693136).

### Sequence mapping, filtering and Q.C

AdapterRemoval (adapterremoval 2.3.1; Schubert et al. 2016) was used to remove adapters and trim both Ns and low-quality bases from the read-ends. Trimmed sequencing data were aligned against the reference genome, gasAcu1-4 (Roberts Kingman et al. 2020), using the Burrows-Wheeler Alignment Tool (BWA, version 0.7.17-r1198-dirty) with the aln algorithm (Li and Durbin 2009), disabling seeding (option -l 1024) thereby increasing mapped data by including reads with post-mortem damage at the read-ends (Schubert et al. 2014). The resulting bam files were sorted and merged using Samtools (version 1.16.1; Li et al. 2009). Duplicate reads in all sorted bam files were identified by Samtools markdup. Masked regions encompassing interspersed repeats and low complexity DNA sequences detected by Repeat-Masker (Smit et al. 2015) and covering ∼4 % of the stickleback genome, and highly repetitive DNA sequences detected by WindowMasker (Morgulis et al. 2006) from the NCBI C++ toolkit covering ∼26 % of the stickleback genome (constructed using -sdust true as setting) were removed from the bam files using Bedtools (version 2.26.0; Quinlan 2014). The total number of reads, average read length, mapping quality and the average depth of coverage on the marine-freshwater divergent sites after the mapping and filtering of the total sequencing reads for each ancient stickleback genomes are listed in the Supplementary Table 3.

### Assessing postmortem DNA damage and contamination

Analyses of potential nucleotide misincorporations using PMDtools (Skoglund et al. 2014) to compare with the modern reference genome revealed that sequencing reads exhibited various levels of characteristic post-mortem damage patterns (Supplementary Figure 2; Dabney et al. 2013; Briggs et al. 2007). Ancient threespine stickleback single-strand libraries exhibited C > T changes at both the 5’ and 3’ read termini. In contrast, double-strand libraries exhibited an excess of G > A (the reverse complement of C > T) changes at the 3’ read termini.

### Metagenomic analyses

We investigated the metagenomic content of our aDNA libraries by competitively mapping our sequencing data, using the pipeline described above, against the RefSeq mitochondrial database v.92 (a concatenated set of eukaryote mitochondrial genome sequences in fasta format). The threespine stickleback mitochondrial reference (NC_041244.1) was the mitogenome with highest number of mapped reads for the ancient samples from Jossavannet, Klubbvatnet, Lilletjønn, Ravnsborgtjernet, and Digerudtjern (Supplementary Table 4). The Borrevannet and Hoksavannet samples were not included in the metagenomic analyses as we only had post-capture sequence data and no shotgun sequence data for these samples.

### Modern stickleback genome reference panel

A dataset of 553 contemporary marine and freshwater threespine sticklebacks was constructed as a reference ancestry panel. To include contemporary sticklebacks descended from recent shared ancestors with the ancient genomes, we returned to lakes and adjacent fjords where the ancient samples had been found in the sediment cores (Figure 1A & B). However, at several of these lakes we failed to capture any sticklebacks in the traps after 2-3 days, suggesting that they no longer supported contemporary stickleback populations. Some had transitioned from lakes to marshland due to gradual infill of organic material (peat), or contained known stickleback predators such as dragon fly larva (*Aeshna sp*.) and eels (*Anguilla anguilla*). See Supplementary Table 2 for the final sample size of contemporary stickleback from each lake. DNA was extracted from fin clips using Qiagen’s DNeasy Blood and Tissue Kit (ID:69581). Sequencing libraries were built using the NEBNext® Ultra™ II FS DNA Library Prep Kit for Illumina (E7805L) in the laboratory facilities of UiO and NTNU, Norway. These libraries were sequenced 150bp PE across a lane of an Illumina NovaSeq 6000 S4 flow cell by the Norwegian Sequencing Centre.

We further added sequencing data available in open repositories from populations across the Northern hemisphere range of the species (Figure 1A) resulting in total dataset of 553 individuals (Supplementary Table 2). The Roberts Kingman et al. (2021) dataset included genomes of threespine stickleback from Loberg lake, which are freshwater individuals from a population that was established sometime between 1983-1988 by anadromous (marine) threespine sticklebacks, following their earlier eradication with rotenone treatment in 1982. Thus, their genomes should reflect the early stages of freshwater adaptation (<30 generations). Sequencing data for all genomes were aligned against the reference genome gasAcu1-4 (Roberts Kingman et al. 2020; doi:10.5061/dryad.547d7wm6t) and processed the same as the data from the ancient genomes with the exception of using Burrows-Wheeler Alignment Tool (BWA, version 0.7.17-r1198-dirty) with the mem algorithm (Li and Durbin 2009) for the contemporary genomes.

### Principal Component Analysis

We used ANGSD 0.940 (Korneliussen et al. 2014) to call pseudo-haploid genotypes of contemporary and ancient genomes, and to then estimate covariance within freshwater-marine divergence associated regions (Jones et al 2012) among genomes. The ancient genomes were included in the principal components computations and not projected onto principal components of modern genomes. Our approach can provide an indication of whether the ancient genomes were impacted by sequencing- or sequence data processing errors. If this was the case, the ancient genomes could appear as outliers in the PCA. Due to the lower coverage of some of the ancient genomes, missing values were imputed using R package missMDA (Josse and Husson, 2016). To remove the potential bias resulting from the ancient DNA damage patterns, 5 nucleotides were trimmed from both ends of the reads when creating the covariance matrix (*i*.*e*. post-mapping). Additional filtering steps included in these analyses were the removal of regions of poor mapping quality (Q < 20), removal of sites with low base quality scores (q < 20), calling only SNPs inferred with a likelihood ratio test (LRT) of P < 0.000002, a minimum allele frequency of 0.25 and specifying uniquely mapping reads only. After these filtering steps, a total of 50,476 SNP sites were used to estimate the covariance across all genomes in the dataset. The number of SNPs covered in each ancient genome is listed in the Supplementary Table 1. The eigenvectors from the covariance matrix were generated with the R function ‘‘eigen’’. Density plots for the PC1-axis were added to better visualize the segregation between the different marine-freshwater populations. Comparing PCA plots including and excluding ancient samples indicated the ancient samples did not impact the PCA among contemporary samples (Supplementary Figure 3). Four focal genomes were then selected for downstream analyses based upon their distribution along PC1, as we wanted to compare genomes that spanned from being mainly marine-adapted to mainly freshwater-adapted. We then selected the genomes with the highest coverage that achieved the first criteria. These genomic libraries were also identified by the metagenomic analyses as having a high proportion of threespine stickleback DNA sequences, with the exception of the Borrevannet genome.

### Sexing

Ancient samples were sexed using the approach of Kirch et al. (2021). Sticklebacks have an XY sex-determination system, where males are the heterogametic sex. Males are therefore haploid for the X chromosome and diploid for the autosomes, while females are diploid for both the X chromosome and autosomes. The sex of the ancient samples was determined by comparing the mean coverage at an autosome (chrXX), part of the pseudoautosomal region (PAR), and the sex-determining region (SDR) of the X-chromosome (chrXIX). For this we used data mapped to the v5 assembly (Peichel et al. 2020) as the PAR and SDR of the X-chromosome are well-defined in this assembly (see Figure S5 of Peichel et al. 2020). We used the samtools coverage command to estimate the mean depth of coverage between coordinates (chrXIX:1-2000000) and (chrXIX:5000000-20000000), corresponding to the PAR and SDR. Ancient genomes were determined to be male if the SDR had half the mean coverage of the autosome and PAR, and female if all three markers had approximately even coverage. The Lilletjønn sample was found to be a male (SDR/PAR coverage = 0.63) and Ravnsborgtjernet and Jossavannet samples were female (SDR/PAR coverage of 0.93 and 0.99 respectively). This determination for Jossavannet is the same as previously made by Kirch et al. (2021) with lower cover data from double-strand library sequences.

### Freshwater ancestry probability estimation

To investigate the marine and/or freshwater origin of adaptive alleles carried by ancient genomes we assigned the ancestral state of each allele, based on allele frequencies in present-day marine and freshwater populations in our modern dataset of 553 individuals (Supplementary Table 2) using a Bayesian estimator of freshwater ancestry (Kirch et al. 2021; and Laine et al. 2024). We restricted our analyses to sites targeted by the enrichment capture probes and estimated genotype probability at the per-site level. The capture probes are designed to target transversion sites, but transitions were not specifically excluded. In total there were 170,567 informative SNP sites across all pseudo-haploid genomes in the dataset, and we focused our subsequent analyses on those sites that were covered in at least by one ancient genome. The number of sites covered in each ancient genome are listed in Supplementary Table 1.

We defined two possible ancestries for the ancient genome, *A*_*marine*_ and *A*_*fresh*_. The allele detected at sites within the marine-freshwater divergent regions (as identified in Jones et al. 2012) were then compared to the contemporary marine and freshwater stickleback in our reference panels. The probability of observing an allele (0,1) in the ancient genome given a specific ancestral state were calculated as:

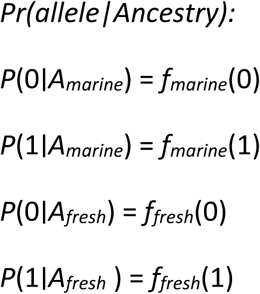

We assume uniform prior probabilities *P(A*_*marine)*_ and *P(A*_*fresh*_), and calculate the posterior probability of each possible ancestral state as:

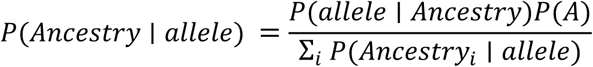

Our estimates are presented as the probability of the allele found at a site in the ancient stickleback being freshwater-associated, with a range of 0 to 1. For P(freshwater ancestry) to be 0 or 1, different alleles should be fixed in the marine and freshwater ecotype genomes within the reference panel. Values greater than 0 but less than 1 indicate the per-site differentiation in (non-fixed difference) allele frequencies between the marine and freshwater ecotype genomes within the reference panel. Thus, a value of 0.5 would suggest that allelic variation at that site was not strongly associated with marine-freshwater divergence. The ancestry probability estimates are therefore not independent from *F*_ST_.

### *F*_ST_ *estimation*

Estimation of *F*_ST_ between two reference set of genomes (one reference set of marine fish and one of freshwater) were performed with vcftools function suite (version 0.1.17; Danecek et al. 2011) from an input file in the variant call format (VCF) created with bcftools mpileup and bcftools call (version 1.5; Li et al. 2009). The marine reference set comprised the ten most marine individuals along the PC1-axis (Figure 2A), and the freshwater reference set comprised the ten most freshwater individuals from the six Norwegian freshwater lake populations included in the modern stickleback dataset. The Norwegian populations were used, as they likely originate from the same ancestral marine population as the ancient samples used in this study, thus providing the most meaningful comparison point.

**Figure 2.**
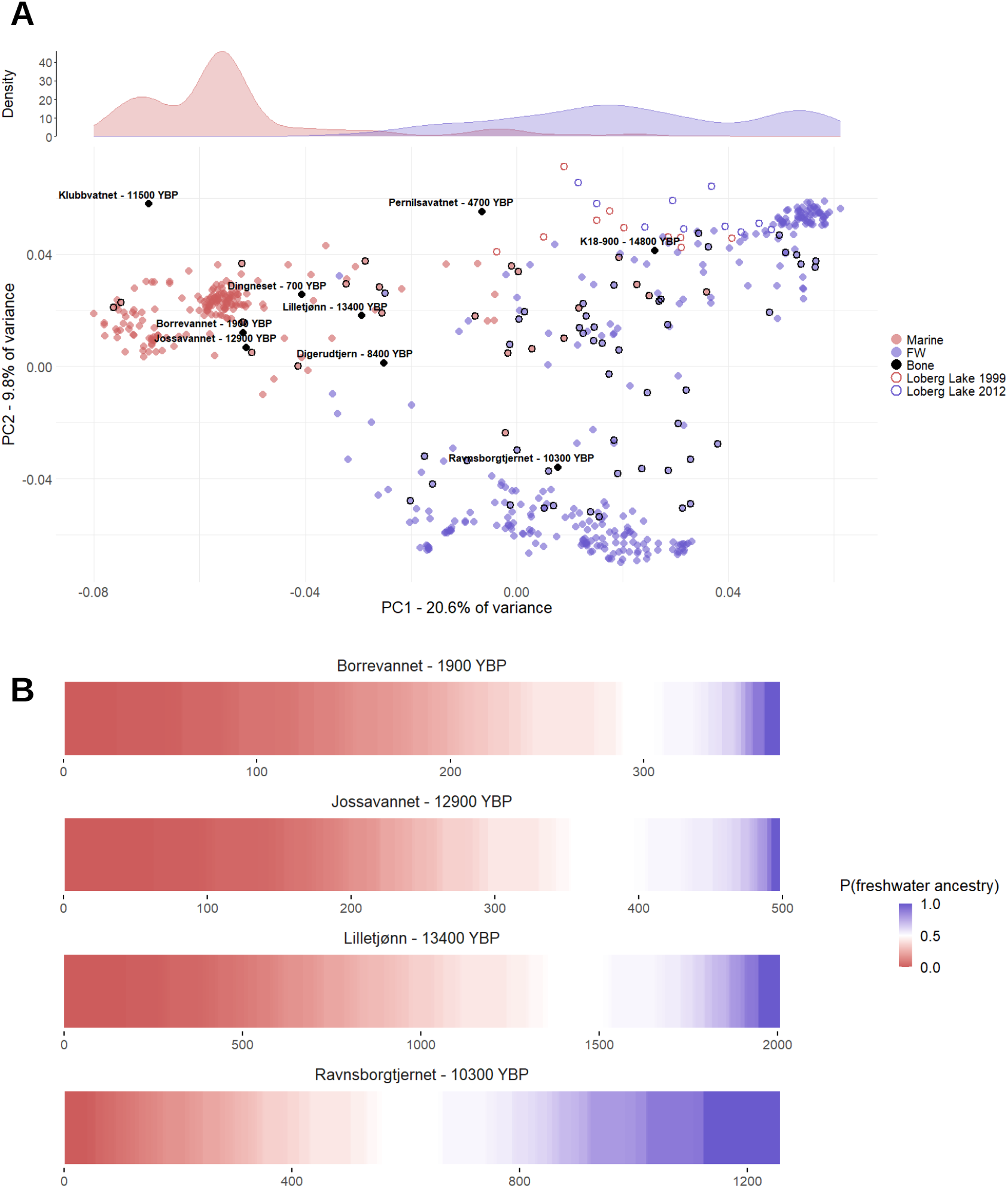
**A**. Principal Component Analysis (PCA) of the marine-freshwater divergent regions of threespine stickleback showing the first and second PCs. The proportion of genetic variance captured by each component is indicated by the axis labels. Ancient genomes are represented by the black markers, contemporary marine stickleback by red markers and contemporary freshwater stickleback by blue markers. Individual stickleback were assigned as marine or freshwater based on sampling location. The density distribution plot above the PCA shows the density of individuals assigned to marine or freshwater populations along PC1. Genomes representing the experimental release populations from Loberg Lake, Alaska are indicated by open markers (Roberts Kingman et al. 2021). **B**. The per-site probability of freshwater ancestry for the marine-freshwater divergent regions of the genome in the four highest coverage threespine stickleback paleogenomes. Sites are ordered based on probabilities from 0 to 1. The probability of freshwater ancestry is dependent upon the allele sampled from the paleogenome, and the allele frequencies sampled from the contemporary freshwater and marine populations. For probabilities to be either 0 or 1, alternative alleles must be fixed in the marine and freshwater ecotype.

The reference stickleback genome gasAcu1-4 (Roberts Kingman et al. 2020) and the twenty modern genomes were used as input for bcftools mpileup (version 1.5; Li et al. 2009). The output file was subsequently piped into bcftools call which used the multiallelic-caller to output a VCF file for all twenty modern genomes (Li et al. 2009). The VCF file was filtered with vcftools (version 0.1.17; Danecek et al. 2011) for sites with a minor allele frequency greater than or equal to 0.05, at least 90% of the data non-missing, a quality value above 30, and mean depth values as well as genotypes with a depth of 5. Thereafter, the resulting VCF file was used for calculating *F*_ST_ estimate per site between the two populations based on Weir and Cockerham’s method (Weir and Cockerham 1984) by using vcftools (version 0.1.17; Danecek et al. 2011). The *F*_ST_ estimate was processed and plotted in R (R Core Team. 2021). We estimate *F*_ST_ per site due to the fine-scale variation in marine-freshwater divergent ancestry, *e*.*g* at the 16-kb minimal haplotype at the *EDA* locus (Colosimo et al. 2005; Jones et al. 2012).

## Results and Discussion

Marine and freshwater ecotypes of threespine stickleback are an exemplar study system of parallel adaptation from standing genetic variation (Reid et al. 2021). By incorporating ancient stickleback genomes sampled from the transitionary layers of lakes formed since the last ice age, we were able to shed new light on the tempo and chronology of genomic changes associated with freshwater adaptation.

### The tempo of freshwater adaptation

Contemporary sticklebacks segregate into marine and freshwater clusters along PC1 of a PCA with overlap in the tails of the density distribution for each ecotype (Figure 2A). Freshwater individuals formed two clusters along PC2, one mainly encompassing the Pacific freshwater populations, the other the Norwegian freshwater populations. The Pacific freshwater cluster separates further than the Norwegian freshwater cluster from the marine cluster along PC1, possibly reflecting the larger pool of freshwater adaptive alleles present in the Pacific stickleback populations (Roberts Kingman et al. 2021). Individuals in the centre of the PC1 axis represented the genotypes of stickleback sampled from rivers (e.g. Big River, California; Little Campbell River, British Columbia; and the Nidelva River, Trondheim). Populations of threespine stickleback in rivers can have a cline from freshwater to increasing marine ancestry towards where the river mouth meets the sea (e.g. Jones et al. 2012).

The ancient genomes span from the marine cluster to the freshwater clusters along PC1. This distribution potentially reflects that the ancient genomes represent different stages of freshwater adaptation. However, the low coverage of some ancient genomes could bias the position of these data points in the PCA. Therefore, we focused on the four ancient genomes with the highest coverage in downstream analyses. Ancient genomes Borrevannet, Jossavannet, Lilletjønn and Ravnsborgtjernet had coverage at 377, 500, 2,035 and 1,279 sites respectively in the marine-freshwater divergent regions of the genome (Figure 2B). Estimating the per-site probability of freshwater ancestry for these four ancient genomes confirmed their position in the PCA plot (Figure 2B). Borrevannet and Jossavannet genomes, and to a lesser extent the Lilletjønn genome, contained mostly marine ancestry, suggesting these two stickleback populations were at the early stages of freshwater adaptation. In contrast, the Ravnsborgtjernet genome carried a marginally higher proportion of freshwater than marine alleles reflecting an intermediate stage of the freshwater adaptation, (Figure 2B).

The Ravnsborgtjernet stickleback subfossil was recovered from a layer dated 10.3 KYR BP (Figure 1C; Supplementary Table 1). At that time and until at least 10.0 KYR BP, the fjord environment was dominated by meltwater from the decaying ice sheet in the interior of southern Norway (Romundset et al., 2023; Romundset et al., submitted.), which likely affected the stratification and salinity conditions, of the fjord waters (Sejr et al. 2022). Reduced salinity at the time could explain the increase in frequencies of the alleles beneficial in the freshwater environments in the early post-glacial Oslofjord population. Such a correlation between salinity gradients and freshwater genotypes is found within the present-day Baltic Sea (DeFaveri et al. 2013; Guo et al. 2015). This hypothesised increase in freshwater standing genetic variation in the founding Oslofjord population could underpin the higher proportion of freshwater alleles at marine-freshwater divergent sites in the Ravnsborgtjernet ancient genome compared to our other ancient stickleback genomes recovered from corresponding transitional sediment layers.

A further consideration is the caveat that ancient genomes represent a single individual and may not be representative of the ancestral population allele frequencies of each lake at that time. As an example, the Jossavannet genome recovered from a bony armour plate and a spine has a higher proportion of marine ancestry than was found in DNA from sediment layers from Jossavannet deposited during and immediately after the transition from a marine to a freshwater habitat (Laine et al. 2024; Supplementary Figure 4). The Jossavannet bones originate from layers when marine sticklebacks could potentially still colonise the lake during high tides, and so may represent outliers from the population mean genotype at that timepoint.

### The chronology of freshwater adaptation

Sorting the marine-freshwater divergent regions covered in the ancient genomes by their physical chromosomal location heuristically revealed the clustering of freshwater alleles (Figure 3A). A few regions on a subset of chromosomes harboured most of the freshwater ancestry alleles; a pattern to some extent shared among all four ancient genomes. The greatest accumulation of freshwater alleles was consistently at the X-chromosome (chromosome XIX in the reference assembly), and within low recombination regions harbouring major effect loci (*EDA*) and regions of parallel marine-freshwater divergence (*WNT7B* and *SULT4A*) on chromosome IV (Figure 3B; Supplementary Table 5). This pattern is consistent with expectations of regions with low recombination rate (Roesti et al. 2013; Samuk et al. 2017; Venu et al. 2024), which reduces the crossover events and disassociation of aggregated haploblocks beneficial in the freshwater environment. In the case of the X-chromosome, selection can more directly act on recessive freshwater-adaptive genetic variation when carried in the male individuals. Aligning the marine-freshwater divergent regions covered in the ancient genomes with per-site *F*_ST_ values between contemporary marine and freshwater sticklebacks at the same sites revealed which regions of high differentiation (*F*_st_) showed freshwater ancestry (Figure 3B). When considering all regions of high *F*_ST_ between marine and freshwater stickleback we find three of the four ancient sticklebacks (excluding Ravnsborgtjernet) still have marine ancestry at high *F*_ST_ sites in the ancient sticklebacks (Figure 4A). Therefore, based on our understanding of the genetic basis of parallel freshwater adaptation from contemporary sticklebacks (e.g. Jones et al. 2012; Roberts Kingman et al. 2021), freshwater adaptation is incomplete in these particular ancient sticklebacks. In contrast, high *F*_ST_ sites on chromosome IV have freshwater ancestry in the Ravnsborgtjernet, Borrevannet and Lilletjønn genomes (Figure 4B). We can therefore predict from these freshwater genotypes at marine-divergent regions on chrIV, that these three ancient sticklebacks would have shared the freshwater phenotype encoded by chrIV loci, such as *EDA*. We suggest this result reflects that these phenotypic traits are under strong selection and are therefore some of the first to show genetic signatures of freshwater adaptation. This same overall pattern in the freshwater ancestry was observed in the recent release-recapture study by Roberts Kingman et al. (2021) suggesting a potential pattern of parallel chronology of genetic changes associated with freshwater adaptation across threespine stickleback populations.

**Figure 3.**
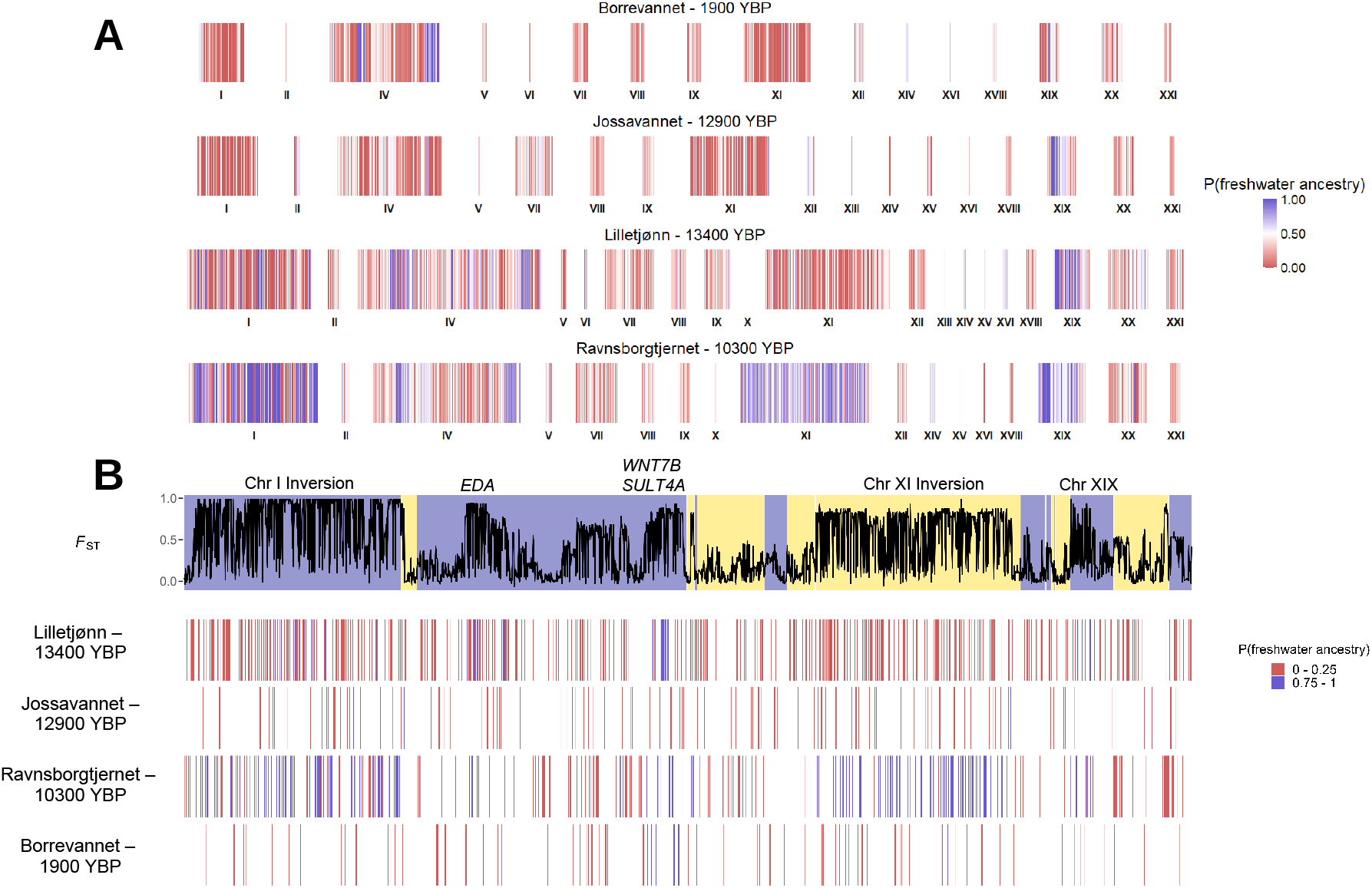
**A**. Per-site probability of freshwater ancestry for the marine-freshwater divergent regions of the genome from Figure 2B, with sites ordered based on their physical position in the genome. Sites are clustered based on the chromosome, and chromosome identities are indicated with Roman numerals under each cluster. Chromosomes III and XVII are omitted from the plot as none of the four high coverage paleogenomes had freshwater ancestry probability values calculated on any site on these chromosomes. The probability of freshwater ancestry is dependent upon the allele sampled from the paleogenome, and the allele frequencies sampled from the contemporary freshwater and marine populations. For probabilities to be either 0 or 1, alternative alleles must be fixed in the marine and freshwater ecotype. **B**. *F*_ST_ values of modern marine and freshwater threespine stickleback populations on the marine-freshwater divergent regions plotted with per-site probability of freshwater ancestry in the four highest coverage paleogenomes on the corresponding sites. The sites are ordered based on the physical chromosomal position in the genome. Only the highest and lowest quartiles of the freshwater ancestry probability value sites are presented. Sites in different chromosomes are indicated with the alternating background colour of the *F*_ST_ value line graph. Sites with high *F*_ST_ value in the modern stickleback populations coinciding with high freshwater ancestry likelihood in the paleogenomes located on inversions with reduced recombination rate, sex chromosome (chromosome XIX) and major effect loci associated with threespine stickleback freshwater adaptation are annotated above the *F*_ST_ value line graph.

**Figure 4.**
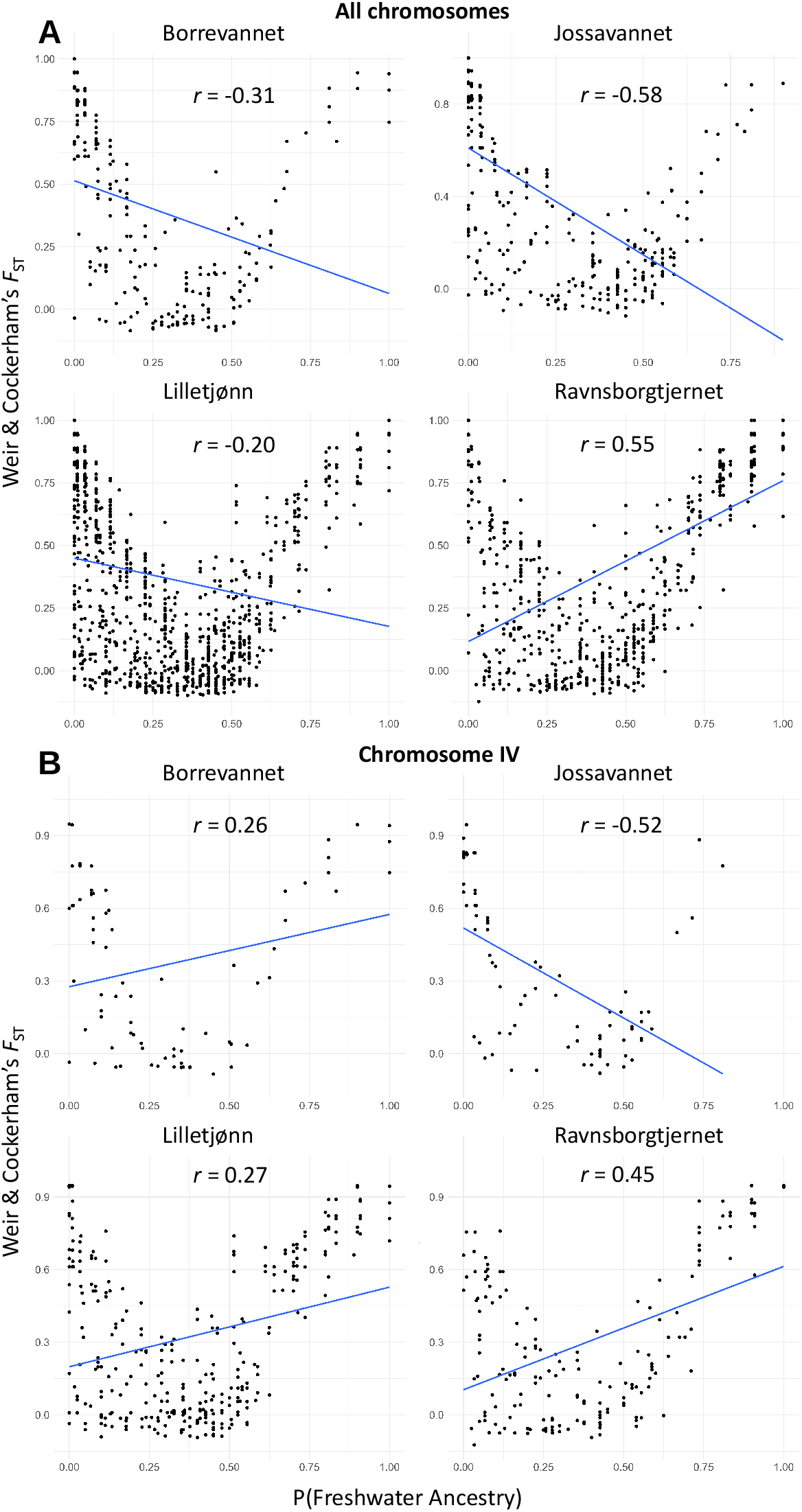
**A**. Per-site *F*_ST_ value in the modern threespine stickleback populations plotted against freshwater ancestry likelihood in the four high coverage paleogenomes on the marine-freshwater divergent region sites covered in each paleogenome across all the marine-freshwater divergent regions of the genome, and **B**. across the marine-freshwater divergent regions of chromosome IV. The blue line in each sub-panel visualizes the linear relationship between the *F*_ST_ value and the freshwater ancestry likelihood, and the *r-*value shows Pearson’s correlation between the two variables. Note that *F*_ST_ are non-indepentent variables as ancestry scores are dependent upon the differentiation between marine and freshwater stickleback reference panels. However, whether the allele in the ancient genomes at high *F*_ST_ sites reflects major allele in marine or freshwater reference panel is of interest here.

In the Ravnsborgtjernet genome we find that further freshwater adaptive variation has accumulated in known inversions in chromosomes I and XI (Figure 3). Inversions have been shown to disproportionately contribute to parallel changes in ecotypes of other systems (e.g. dune sunflowers *Helianthus petiolaris*, Huang et al. 2025, seahorses (*Hippocampus guttulatus*, Meyer et al. 2024, and marine snails *Littorina saxatilis* Koch et al. 2022). Inversions are typically characterised by low recombination, and can thereby retain linkage between multiple adaptive loci, a trait that has been proposed to explain their disproportionate use in parallel evolution (Westram et al. 2022).

## Conclusions

Our analyses of ancient threespine stickleback genomes across different sites deepens our understanding of the early-stage freshwater adaptation in threespine stickleback. We find regions of reduced recombination containing multiple large effect loci in the threespine stickleback genome are consistently among the first to harbour freshwater alleles within the marine-freshwater regions of the genome. The genetic architecture of the regions repeatedly showing the initial stages of parallel freshwater adaptation in threespine stickleback is characteristic of the genomic architecture broadly predicted to facilitate parallel evolution (Orr 2005; Bailey et all. 2017; MacPherson & Nuismer 2017). As such our findings may have broader significance for understanding the chronology of genetic changes underpinning parallel adaptation in nature.

## Acknowledgements

We thank two anonymous reviewers for their feedback on an earlier draft of this manuscript, and Jacob Enk for helpful feedback on Mybaits capture protocol. Martin Buran and Lina Gislefoss of NGU helped with coring fieldwork. Mike Martin (all Holomuseomics) and Sarah Martin provided useful discussions and advice regarding lab work. We additionally thank Tom Gilbert, Nuno Martins and Sarak Mak for help with earlier lab work using facilities in Copenhagen. This work was funded by an ERC Consolidator Grant – EXPLOAD (101045346) awarded to ADF.

