## Supplementary Figures for "Ancient stickleback genomes reveal the chronology of parallel adaptation"

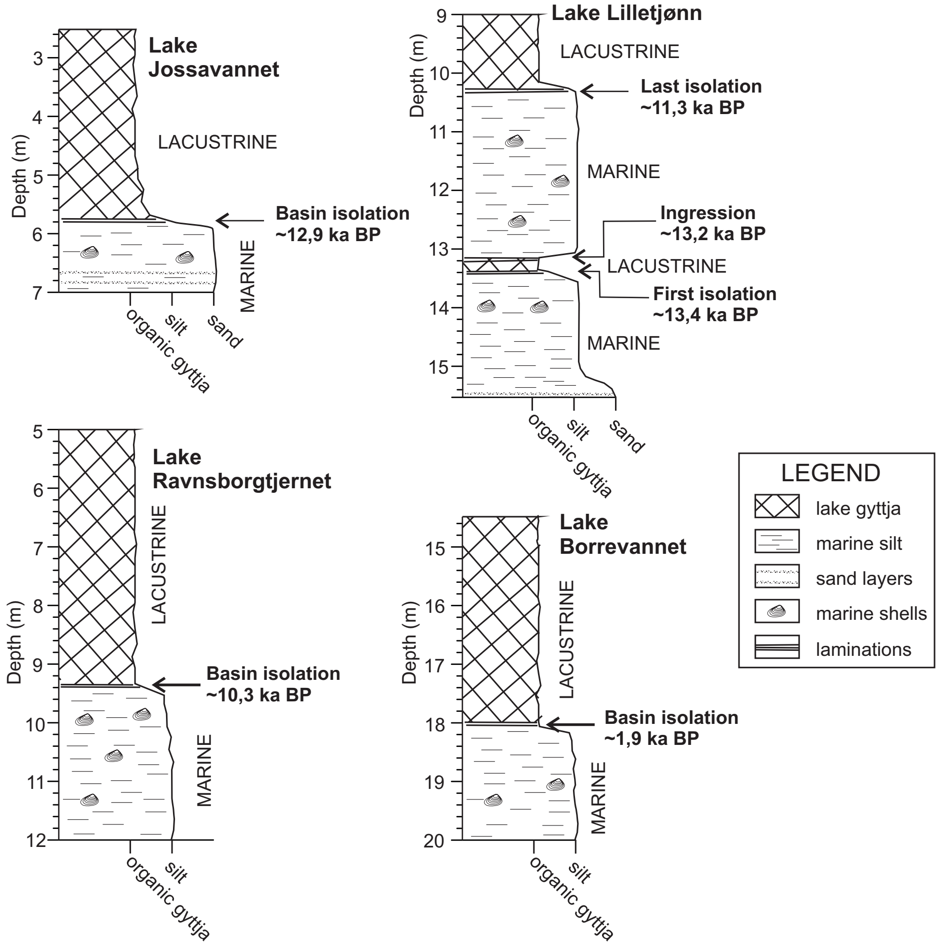


**Supplementary Figure 1.** Schematic diagrams of the stratigraphy in the study lakes from which the four high coverage paleogenomes were recovered. Lake Lilletjønn in this figure corresponds to the Kristiansand sample, as this is the official name of this lake from which these subfossil remains were recovered.

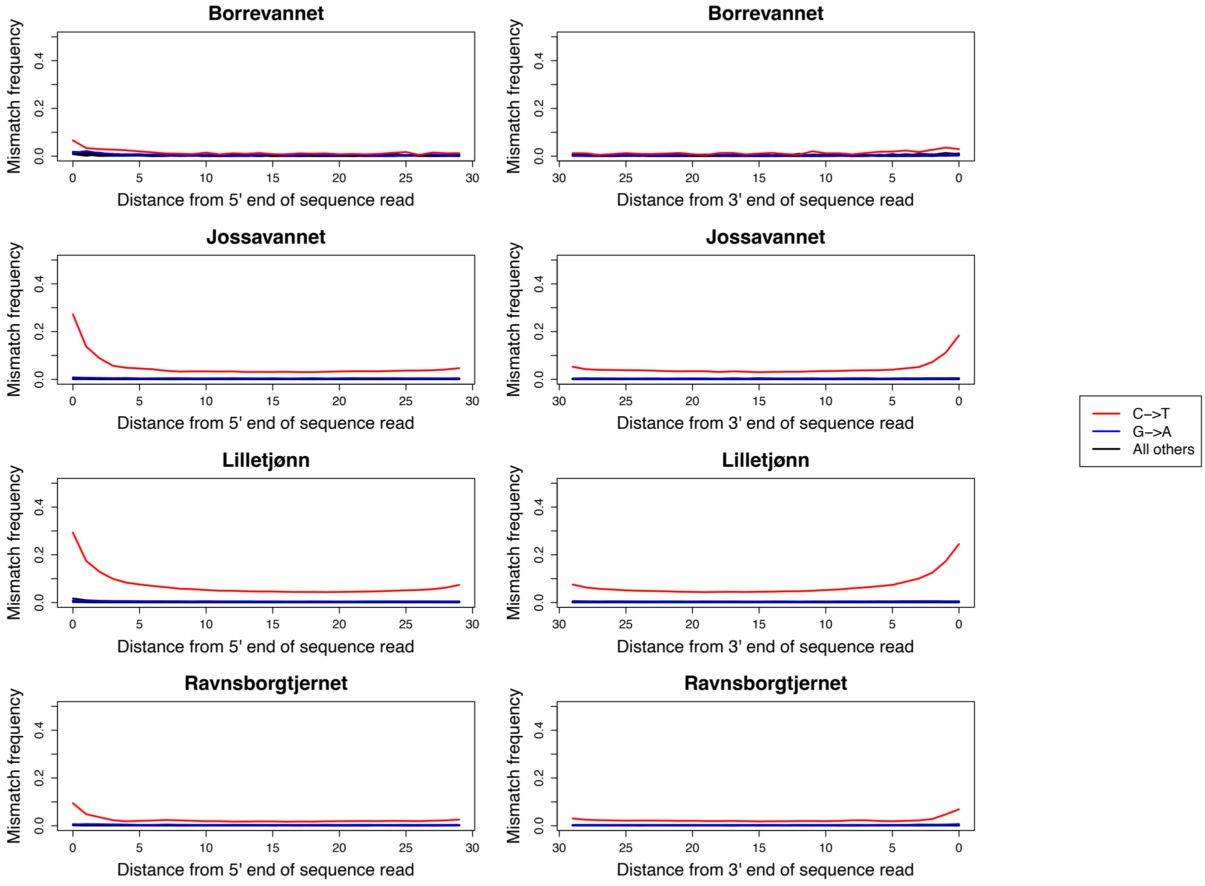


**Supplementary Figure 2.** Post-mortem deamination patterns for filtered reads sequenced from stickleback bones, armour plates and spines and mapped against the three-spined stickleback reference genome gasAcu1-4. Plots show an excess of C→T sequence misincorporation errors at the 5′ and 3′ read termini relative to the modern stickleback reference characteristic of DNA damage in single-strand library builds on ancient DNA sample. The extent of the damage in these four samples corresponds with their relative age, see Supplementary Table 1.

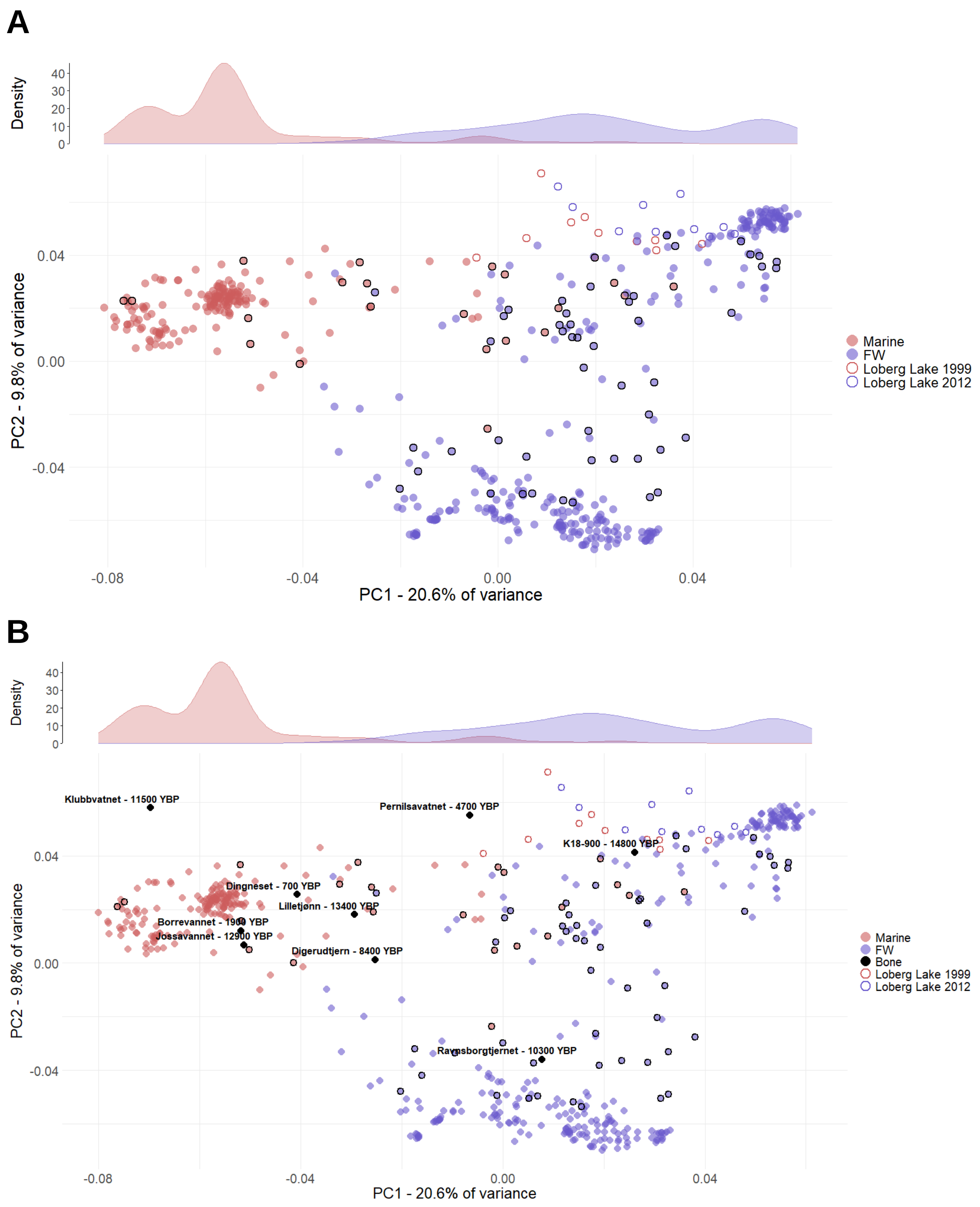


**Supplementary Figure 3.** A comparison of Principal Component Analysis (PCA) plots of the marine-freshwater divergent regions of threespine stickleback, **A.** excluding and **B.** including the ancient samples.

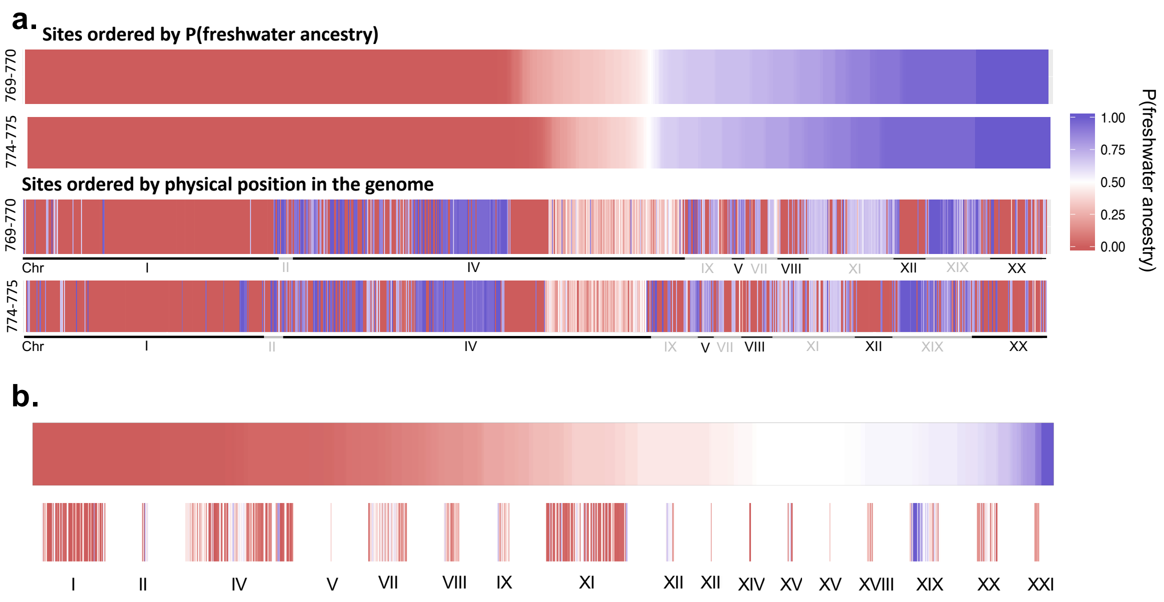


**Supplementary Figure 4.** A comparison of the ancestry in marine-freshwater regions of the stickleback genome using DNA extracted from **a.** ancient sediments from Jossavannet corresponding to a layer midway through the transition from marine to freshwater (774-775 cm depth), e.g. when marine water could still ingress the lake at high tides; and the first freshwater layer (769-770 cm depth) following complete isolation from marine ingression. **b.** DNA extracted from the subfossil hard parts on an individual stickleback. The fossil and sediment layers are dated to 12.9 ka BP (Kirch et al. 2021; Laine et al. 2024). The figure panels show individual transversions shaded on a scale of 0-1 of the probability that the allele at that site is freshwater associated (conditioned on allele frequencies in the contemporary marine and freshwater reference panels). Sites are alternatively ordered by P(freshwater ancestry) and physical position in the genome.
